# Moderate prenatal alcohol exposure differentially alters acute ethanol sensitivity of GABAergic transmission in CRFR1- and CRFR1+ CeM Neurons

**DOI:** 10.64898/2025.12.12.693987

**Authors:** Sarah E. Winchester, Marvin R. Diaz

## Abstract

**Abstract:** Prenatal exposure to alcohol (PAE) increases the risk for misusing alcohol and/or developing an alcohol use disorder (AUD) by adulthood. The corticotropin releasing factor (CRF) system is a major target of pre- and post-natal ethanol (EtOH) exposure. CRF and its receptor (CRFR1), in part, mediate EtOH potentiated GABA release in the medial nucleus of the central amygdala (CeM) of adult male rodents. Interestingly, our lab has shown a disruption in the function and expression of CeM CRFR1 and acute EtOH’s effects on GABA transmission in PAE adolescent animals, but it is unknown whether these alterations to the CRF system persist into adulthood or alter the actions of acute EtOH on GABAergic transmission in the CeM. Using CRF_1_-Cre-^td^Tomato rats, this study examined how *moderate* PAE alters acute EtOH modulation of GABAergic neurotransmission onto CRFR1+ and CRFR1-CeM neurons in adult offspring (P80-105). Pregnant dams were exposed to vaporized ethanol or room air (control) on gestational day 12 (G12) for 6 hours and whole-cell electrophysiology was performed in the CeM to assess the actions of acute EtOH (44, 66, & 88 mM) on GABAergic transmission onto CRFR1+ and CRFR1-neurons. We found unique effects of PAE that were cell type- and concentration-dependent in males and females, suggesting PAE dysregulates acute EtOH’s modulation of GABA transmission within the CeM in a sex-specific manner. This study contributes to the expanding body of research exploring the effects of PAE and how a single exposure can impact neurophysiological mechanisms in brain regions associated with AUD.

**Highlights:**

- *Moderate* PAE alters the actions of alcohol on synaptic transmission in adult rats
- PAE differentially impacts GABAergic transmission onto CeM CRFR1- and CRFR1+
- Long-term PAE effects in the CeM are sex-specific

## 1.0 Introduction

Although public health efforts have increased awareness, alcohol consumption during pregnancy remains relatively common. In the United States, ∼14% of pregnant women reported alcohol use in the last 30 days, with higher prevalence rates observed in several European nations (∼20-28%) (Gosdin et al., 2022; Mardby et al., 2017). Consequently, prenatal alcohol exposure (PAE) can result in Fetal Alcohol Spectrum Disorder (FASD) which can significantly increase the risk for misusing alcohol and/or developing alcohol use disorder (AUD) later in life (Alati et al., 2006; Baer et al., 1998; Baer et al., 2003). The recruitment of the corticotropin releasing factor (CRF) system in the medial central amygdala (CeM) is a major target of ethanol (EtOH), particularly during the development of AUD (Gilpin, 2012; Gilpin et al., 2015). Importantly, our lab has identified a functional change in the CRF receptor type I (CRFR1) in the CeM of adolescent animals following a *moderate* gestational (G) day 12 (G12) PAE, where CRFR1-agonist effects were either blunted or reversed compared to controls (Rouzer and Diaz, 2022). In conjunction with this, G12 PAE increased and decreased CRFR1 mRNA in females and males, respectively (Rouzer and Diaz, 2022). How these PAE-induced changes may persist into adulthood or alter the effects of future exposure to alcohol on neurotransmission is unknown.

Typically, at the level of the synapses, CRF and/or acute EtOH increases GABA release in the CeM of adult male rodents, which is partially mediated by CRFR1 (Nie et al., 2009). Using adult GFP reporter mice, Agoglia & colleagues (2021) found acute EtOH (44 mM) decreased GABA release onto CeM CRFR1+ cells in adult males, with females following a similar pattern, though nonsignificant (Agoglia et al., 2020). Consistent with these reports, the CeM of female rodents appears to be insensitive to the effects of acute EtOH (Kirson et al., 2021; Winchester and Diaz, 2025). Interestingly, we recently showed that *moderate* G12 PAE differentially affects acute ethanol actions on GABA transmission in a concentration- and sex-specific manner in CeM neurons of adolescent rats (Winchester and Diaz, 2025). However, whether PAE-induced neuroadaptations alter the effects of EtOH on GABA transmission within the CeM in adult rats remains unclear. Using CRF_1_-Cre-^td^Tomato rats (Weera et al., 2022), the current study investigated how *moderate* PAE on G12, an important developmental period for the amygdala (Soma et al., 2009), may alter acute EtOH modulation of GABAergic neurotransmission in CRFR1+ and CRFR1-CeM neurons in adult offspring. We hypothesized that PAE would attenuate or reverse the acute effects of EtOH across all concentrations in PAE males relative to male controls and would elicit opposing acute EtOH effects at the highest concentration (88 mM) in females compared to female controls.

## 2.0 Materials and Methods

### 2.1 Animals and *Moderate* Prenatal Alcohol Exposure (G12 PAE)

Male CRF_1_-Cre transgenic rats were obtained for breeding from Dr. Nicholas Gilpin, Louisiana State University Health Sciences Center (New Orleans, LA, USA). Breeding was done in-house as previously described (Mooney and Varlinskaya, 2011; Rouzer et al., 2017; Rouzer and Diaz, 2022; Winchester and Diaz, 2025). Briefly, Cre^+^ male breeders were paired with 2 wild-type Wistar females obtained from Inotiv (Indianapolis, IN, USA) for 4 days, where vaginal smears were obtained daily for detection of sperm, and detection was classified as gestational day (G) 1. Pregnancy viability was monitored via dam body weight on G1, G10, and G20. On G12, dams were exposed to vaporized ethanol for 6 hr as previously described (Przybysz et al., 2023; Rouzer et al., 2017; Rouzer and Diaz, 2022; Winchester and Diaz, 2025) with a target blood EtOH concentration (BEC) of 60-120 mg/dL. Tail blood collection from EtOH and air-exposed dams occurred at hr 6, and was analyzed using an Analox AM1 Alcohol Analyser (Analox Instruments Ltd, The Vale, London). Upon parturition, pups were sexed and culled to 12 on postnatal day (P) 2, maintaining a 1:1 male to female ratio, when possible, and weighed on P7 & P12. Between P14-P17 pups were ear punched and genotyping for expression of *iCre* was conducted by Transnetyx (Cordova, TN, USA). At P21, pups were weaned and paired-housed with same sex litter mates of the same genotype, when possible. Whole-cell electrophysiology was performed in adult (P80-110) offspring produced from air (n=26) and EtOH (n=25) exposed litters, with no more than 2 animals per sex, per litter included in the study. All animals were housed in a temperature-controlled colony, on a 12h light/dark cycle, with *ad libitum* access to food and water as previously described (Winchester and Diaz, 2025). Binghamton University’s Institutional Animal Care and Use Committee (IACUC) approved all experiments.

### 2.2 Slice electrophysiology

Slice preparation procedures were performed as previously described (Winchester, 2025). Briefly, rats were anesthetized (5% isoflurane), decapitated, brains were quickly removed and placed in oxygenated sucrose artificial cerebral spinal fluid (ACSF). Coronal slices containing the CeM (300 µm) were obtained using a vibratome (Leica Microsystems. Bannocknurn, IL, USA) and incubated for at least 40 min in 33⁰C oxygenated ACSF. Electrophysiology recordings were performed 1-5h following slice preparation.

Experimental procedures and timeline used for whole-cell electrophysiology data collection and analysis were the same as previously described (Winchester and Diaz, 2025). Briefly, upon a successful patch, neurons were allowed to stabilize for 5 minutes. Using a KCl internal solution, spontaneous inhibitory postsynaptic currents (sIPSCs) were recorded to assess the effect of EtOH (44, 66, or 88 mM) on GABAergic transmission onto neurons containing CRFR1 (CRFR1+) and those without (CRFR1-) - recordings were saved for manual analysis using MiniAnalysis (Synaptosoft). In all experiments, each concentration of EtOH was tested in a single cell per animal, with up to 6 separate cells collected per animal. Recordings were included only if access resistance changed by less than 20%.

### 2.3 Statistics

Epidemiological data and animal studies have highlighted sex differences in alcohol consumption (Juarez and Barrios de Tomasi, 1999; White, 2020) and neurophysiological responsiveness to acute EtOH (Agoglia et al., 2020; Agoglia et al., 2022; Kirson et al., 2021), therefore the current study separated the data by sex *a priori* for analysis. Statistical analysis, including tests for normality, t-tests and two-way ANOVAs, were performed using GraphPad 9 software (Prism). Grubb’s test was used to identify a single outlier within each concentration group based on EtOH-induced % change from baseline in sIPSC frequency (Hz). Identified outliers were removed from the corresponding frequency and amplitude % change data, raw sIPSC values, and baseline measurements. All data was tested for normality, and the appropriate parametric test was performed. Significance was defined as p ≤ 0.05, and in the event of a significant interaction, Sidak’s multiple comparison tests were used to determine specific group differences. All data is presented as the mean ± standard error of the mean (SEM).

## 3.0 Results

### 3.1 *Moderate G12* PAE – Blood EtOH Concentrations (BECs), gestational and litter characteristics

Tail bloods were collected from all pregnant rats at the end of the vapor PAE exposure (hr 6). Only litters from pregnant rats that reached BECs of 60-120 mg/dL were used for this study. Specifically, the minimum BEC was 60.40 mg/dL and the maximum 120.1 mg/dL - the average BEC was 90.32 (3.114) mg/dL (**Figure 1**), classifying this exposure as *moderate* as previously described (Valenzuela et al., 2012; Winchester and Diaz, 2025). As previously stated, gestational weights were monitored, and we observed no significant differences between PAE and air exposed dams. Additionally, postnatal weights were not affected by PAE (**Table 1**).

**Figure 1.**
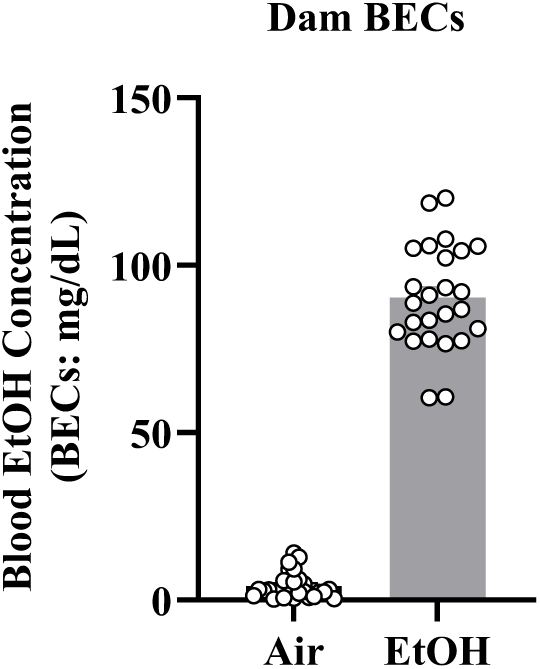
BECs from pregnant dams. (**A**) BECs collected at the conclusion of G12 vaporized EtOH or air exposure. Target BECs = 60-120 mg/dL for EtOH condition.

**Table 1.** Gestational and postnatal litter characteristics, reported as mean (SEM). Gestational and postnatal weights were not altered as a function of PAE. A two-way ANOVA revealed a main effect of day for gestational (F (1.276, 75.31) = 1705, *p <* 0.0001) and postnatal characteristics (F (0.5656, 22.91) = 2344, *p* < 0.0001). * denotes p ≤ 0.05

| **Gestational Day\*** | **<u>Gestational Weights (g)</u>** |  |
| --- | --- | --- |
|  | **Air** | **PAE** |
| G1 | 278.7 (5.560) | 271.4 (3.907) |
| G10 | 308.1 (6.437) | 303.7 (3.967) |
| G20 | 397.5 (6.970) | 389.0 (5.537) |
| **Postnatal Day\*** | **<u>Postnatal Weights (g)</u>** |  |
|  | **Air** | **PAE** |
| P2 | 90.13 (1.834) | 88.08 (2.182) |
| P7 | 183.7 (5.097) | 179.5 (6.199) |
| P12 | 295.6 (6.682) | 292.6 (7.578) |

### 3.2 Electrophysiology

#### 3.2.1 G12 PAE does not alter basal membrane properties

Membrane resistance (Rm) and membrane capacitance (Cm) were monitored throughout recordings to examine the effect of PAE on basal membrane properties. There were no differences observed as a function of exposure in Rm or Cm for CRFR1- or CRFR1+ neurons in males (*p* > 0.05) or females (*p* > 0.05) (**Table 2**). The average access resistance for electrophysiology recordings was 18.86 ± 0.57 for CRFR1-, and 20.78 ± 0.48 for CRFR1+.

**Table 2.** Membrane of CeM CRFR1- and CRFR1+ neurons, reported as mean (SEM). No differences were observed in membrane resistance (Rm) or membrane capacitance (Cm) in CRFR1- or CRFR1+ neurons. A test of normality was conducted before using the proper parametric test.

| **CRFR1-** |  |  |  |  |
| --- | --- | --- | --- | --- |
|  | **Air Males**<br>(n = 32 cells, 20<br>animals) | **PAE Males**<br>(n = 30 cells, 19<br>animals) | **Air Females**<br>(n = 33 cells, 21<br>animals) | **PAE Females**<br>(n = 33 cells, 21<br>animals) |
| **Membrane**<br>**Resistance (MΩ)** | 391.6 (12.06) | 416.0 (18.30) | 443.5 (20.00) | 439.7 (18.23) |
| **Membrane**<br>**Capacitance (pF)** | 48.84 (1.12) | 47.19 (1.27) | 46.02 (1.73) | 47.08 (1.30) |
| **CRFR1+** |  |  |  |  |
|  | **Air Males**<br>(n = 35 cells, 17<br>animals) | **PAE Males**<br>(n = 31 cells, 17<br>animals) | **Air Females**<br>(n = 35 cells, 22<br>animals) | **PAE Females**<br>(n = 31 cells, 18<br>animals) |
| **Membrane**<br>**Resistance (MΩ)** | 418.9 (10.05) | 422.4 (12.91) | 425.6 (12.30) | 422.0 (11.66) |
| **Membrane**<br>**Capacitance (pF)** | 48.79 (1.52) | 46.33 (1.36) | 44.58 (1.39) | 47.23 (1.25) |

#### 3.2.2 G12 PAE does not impact CeM CRFR1- or CRFR1+ neurons under basal conditions

To test the effect of G12 PAE on CeM GABAergic neurotransmission, we assessed the frequency (Hz) and amplitude (pA) of sIPSCs under basal conditions. For CRFR1-neurons (**Figure 2A**), we observed no significant differences in frequency (**Figure 2C**) or amplitude (**Figure 2E**) of sIPSCs in males (*p* > 0.05) or females (*p* > 0.05). Additionally, the frequency (**Figure 2D**) of CRFR1+ sIPSCs was unaffected by PAE in males (*p* > 0.05) and females (*p* > 0.05) (**Figure 2B**). Interestingly, we observed an effect of exposure on the sIPSC amplitude of CRFR1+ (F (1, 127) = 5.026, *p* = 0.027), suggesting PAE produces postsynaptic modifications.

**Figure 2.**
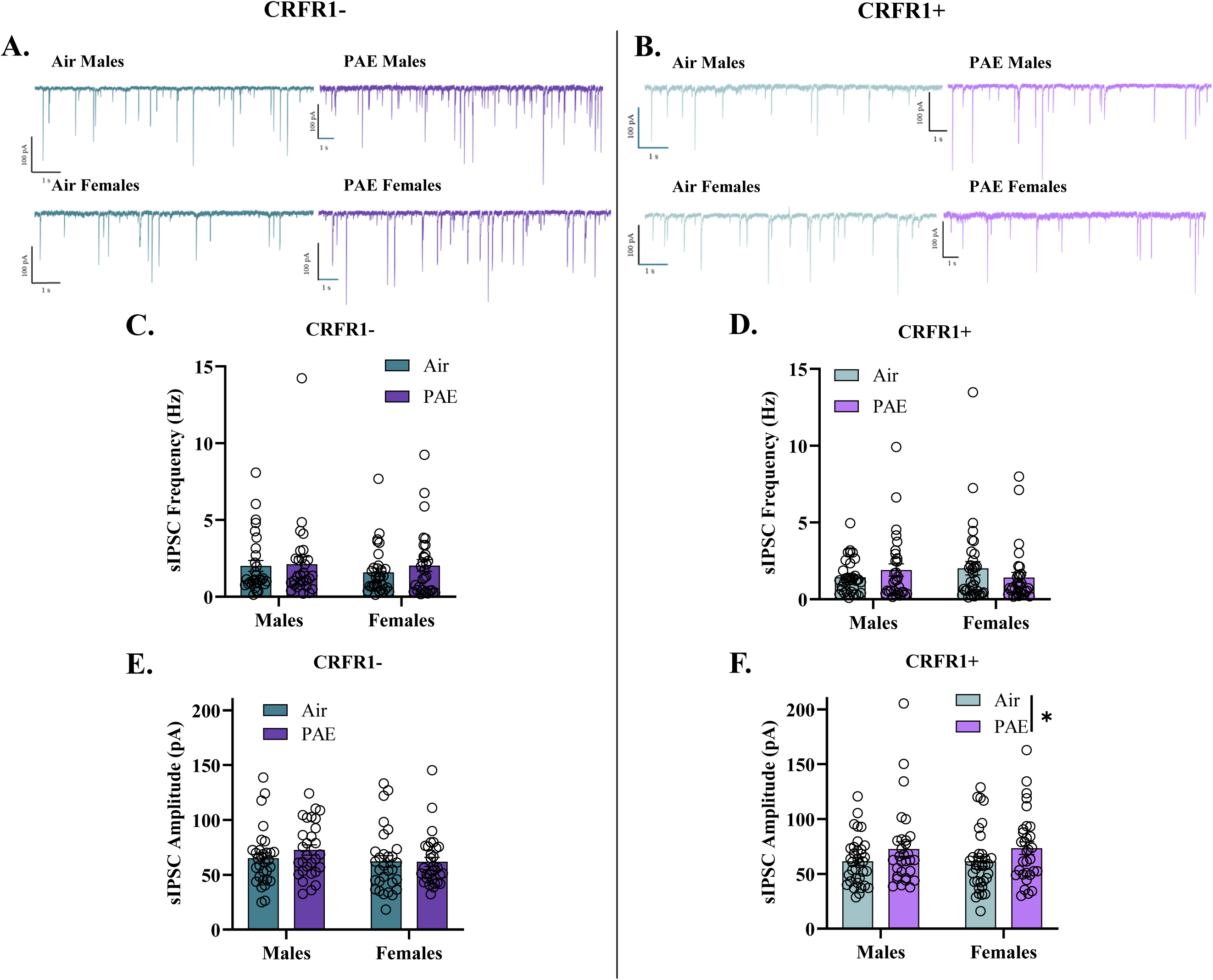
Basal spontaneous inhibitory post-synaptic currents (sIPSCs). Representative sIPSC activity of (**A**) CRFR1-& (**B**) CRFR1+ CeM neurons from males and females. No differences were observed in baseline (**C**) frequency or (**E**) amplitude of CRFR1-neurons (air males: n = 32 cells from 20 animals; PAE males: n = 30 cells from 21 animals; air females: n = 33 cells from 19 animals; PAE females: n = 33 cells from 21 animals). (**D**) CRFR1+ basal frequency was not changed as a function of PAE. (**F**) A main effect of exposure was observed on basal CRFR1+ sIPSC amplitude (air males: n = 35 cells from 17 animals; PAE males: n = 31 cells from 17 animals; air females: n = 38 cells from 22 animals; PAE females: n = 34 cells from 18 animals). PAE = prenatal alcohol exposure. CRFR1 = Corticotropin releasing factor receptor type I. sIPSC = spontaneous inhibitory postsynaptic currents. All data was tested for normal distribution, and the appropriate parametric tests were used. * denotes p ≤ 0.05. Bars represent standard error of mean.

##### G12 PAE males show insensitivity to the actions of acute EtOH in CeM CRFR1-neurons

To examine the effect of G12 PAE on acute EtOH modulation of GABAergic transmission onto CeM CRFR1-neurons, we initially examined raw changes in sIPSC frequency and amplitude (**Supplementary Figure 1**). In air males, 66 mM EtOH decreased the frequency of raw sIPSCs (W = -56.00, *p* = 0.027), suggesting EtOH decreases GABA release onto CRFR1-neurons. In air males, raw sIPSC frequency remained unchanged after application of either 44 mM (t (8) = 0.091, *p* = 0.930) or 88 mM EtOH (W = 11.00, *p* = 0.570). Interestingly, raw sIPSC frequency was not changed by acute EtOH in PAE males regardless of concentration (44 mM: t (9) = 1.232, *p* = 0.249; 66 mM: t (9) = 1.474, *p* = 0.175; 88 mM: W = -11.00, *p* = 0.570; **Supplementary Figure 1A, B, and C**), suggesting an insensitivity to acute EtOH’s effects. Consistent with changes in raw sIPSC frequency, assessment of acute EtOH effects relative to baseline revealed a significant decrease in sIPSC frequency in air males following 66 mM EtOH application (t (11) = 2.658, *p* = 0.022), however this effect was not observed at 44 mM (t (8) = 0.625, *p* = 0.550) or 88 mM (t (8) = 0.602, *p* = 0.564; **Figure 3A**). Additionally, in PAE males, no significant change from baseline was observed in sIPSC frequency following 44 mM (t (9) = 1.151, *p* = 0.279), 66 mM (t (9) = 1.626, *p* = 0.138), or 88 mM (t (8) = 0.733, *p* = 0.484) EtOH application (**Figure 3A**). Finally, G12 PAE had no effect on the dose response curve (F (1, 53) = 0.1291, *p* = 0.721), but we did observe a main effect of concentration (F (2, 53) = 3.132, *p* = 0.051), with no interaction present (F (2, 53) = 0.5618, *p* = 0.574; **Figure 3A**).

**Figure 3.**
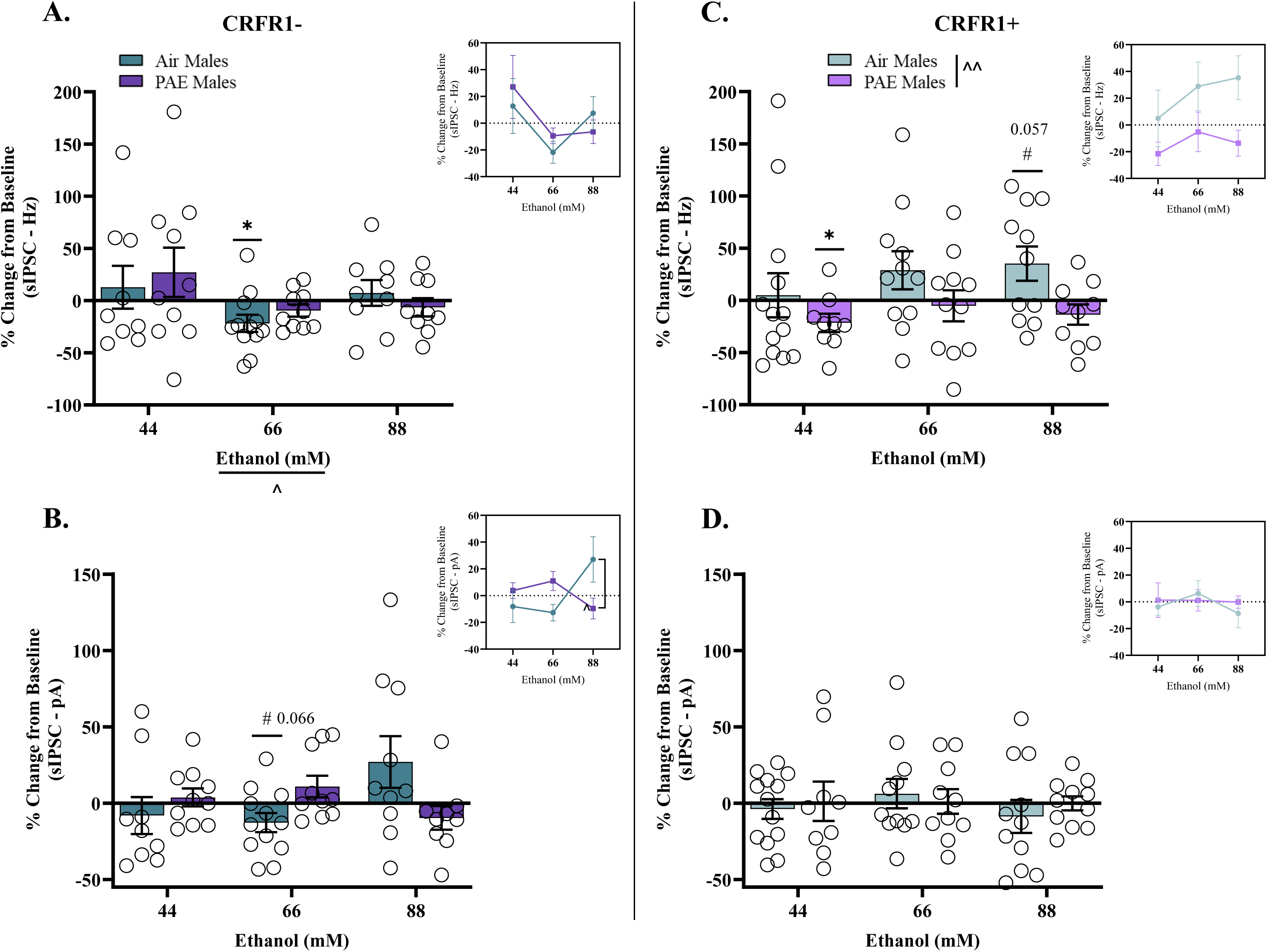
Acute EtOH effects in males represented as % change from baseline. (**A**) A two-way ANOVA revealed a main effect of concentration in CRFR1-neurons. 66 mM significantl decreased sIPSC frequency in air males (n = 12 cells), with no effect observed with 44 mM (n = 9 cells) or 88 mM (n = 9 cells). PAE males were unaffected by acute EtOH at 44 mM (n = 10 cells), 66 mM (n = 10 cells), or 88 mM (n = 9 cells). (**B**) A trend was observed for 66mM to decrease air male sIPSC amplitude of CRFR1-neurons, with no effects observed at other concentrations tested, or in PAE males. (**C**) A main effect of exposure was observed on CRFR1+ frequency. A strong trend was observed for 88mM to increase CRFR1+ air male sIPSC frequency (n = 11 cells), with no effects for either 44 mM (n = 13 cells) or 66 mM (n = 11 cells). 44 mM significantly decreased PAE male sIPSC frequency in CRFR1+ neurons (n = 9 cells), whereas 66 mM (n = 11 cells) or 88 mM (n = 10 cells) had no effect. (**D**) There were no significant differences found on CRFR1+ amplitudes for either group at any concentration tested. PAE = prenatal alcohol exposure. CRFR1 = Corticotropin releasing factor receptor type I. sIPSC = spontaneous inhibitory postsynaptic currents. Inset charts represent a different perspective of the same data set. # denotes trend. ^ denotes significance from two-way ANOVA. * denotes significance from t-test relative to 0. *p* ≤ 0.05. Bars represent standard error of mean.

When assessing sIPSC amplitude in air males, we observed a strong trend for 66 mM EtOH to decrease raw sIPSC amplitudes (t (11) = 2.087, *p* = 0.061), however this effect was not present with 44 mM (t (8) = 1.695, *p* = 0.129) or 88 mM (t (9) = 1.354, *p* = 0.209; **Supplementary Figure 1B, F, and J**). For PAE males, there were no significant changes to raw sIPSC amplitudes at any concentration tested (44 mM: t (9) = 0.109, *p* = 0.916; 66 mM: t (9) = 1.188, *p* = 0.265; 88 mM: t (8) = 1.832, *p* = 0.104; **Supplementary Figure 1B, F, and J**). When analyzing EtOH effect as % change from baseline, we again observed a strong trend for 66 mM EtOH to decrease sIPSC amplitude (t (11) = 2.041, *p* = 0.066; **Figure 3B**), suggesting this concentration (66 mM) may produce postsynaptic modifications in air males. Interestingly, neither 44 mM (t (8) = 0.673, *p* = 0.520) or 88 mM (t (9) = 1.596, *p* = 0.145) produced a significant change from baseline in sIPSC amplitude of air males (**Figure 3B**). Similar to PAE male raw data, we observed no differences from baseline in sIPSC amplitude at any concentration tested (44 mM: t (9) = 0.640, *p* = 0.538; 66 mM: t (9) = 1.534, *p* = 0.159; 88 mM: t (8) = 1.243, *p* = 0.249; **Figure 3B**). Lastly, PAE did not shift the dose response curve of sIPSC amplitudes (F (1, 54) = 0.001963, *p* = 0.965), and there was no main effect of concentration (F (2, 54) = 0.6884, *p* = 0.507). However, a PAE x concentration interaction was observed (F (2, 54) = 5.095, *p* = 0.009); post-hoc analyses revealed an effect of exposure at 88 mM EtOH where air male amplitude increased and PAE male amplitude decreased (*p* = 0.014). Furthermore, the post-hoc analysis revealed a shift in the dose response curve for air males, where amplitudes were decreased at 44 mM (*p* = 0.048) and 66 mM (*p* = 0.013), but increased at 88 mM.

#### 3.2.4 G12 PAE modifies acute EtOH modulation of GABAergic transmission onto CeM CRFR1+ neurons in males

We applied the same statistical approach to assess the effect of G12 PAE on GABAergic transmission onto CRFR1+. We observed no differences in raw sIPSC frequency for air males at any concentration tested (44 mM: W = -35.00, *p* = 0.244; 66 mM: W = 26.00, *p* = 0.278; 88 mM: t (10) = 1.770, *p* = 0.107; **Supplementary Figure 1C, G, and K**). For PAE males, we observed a significant decrease in raw sIPSC frequency at 44 mM (W = -37.00, *p* = 0.027), suggesting a decrease in GABA transmission onto CRFR1+ cells. Conversely, 66 mM (W = -10.00, *p* = 0.700) and 88 mM (W = -27.00, *p* = 0.193) had no effect on raw sIPSCs for PAE males (**Supplementary Figure 1C, G, and K**). When analyzing the data as % change from baseline, we found a strong trend for 88 mM to increase sIPSC frequency in air males (t (10) = 2.147, *p* = 0.057), with no effect observed at either 44 mM (t (12) = 0.233, *p* = 0.820) or 66 mM (t (10) = 1.589, *p* = 0.143; **Figure 3B**).

Interestingly, in PAE males, we observed a significant decrease in sIPSC frequency at 44 mM (t (8) = 2.465, *p* = 0.039), an effect that was not present at either 66 mM (t (10) = 0.350, *p* = 0.734) or 88 mM (t (9) = 1.398, *p* = 0.196; **Figure 3B**). A two-way ANOVA was used to examine shifts in the dose response curve, revealing a main effect of exposure (F (1, 59) = 7.255, *p* = 0.009), with no effect of concentration (F (2, 59) = 0.9363, *p* = 0.398) or an interaction (F (2, 59) = 0.2328, *p* = 0.793).

For air males, we observed no changes in the raw sIPSC amplitude at any concentration tested (44 mM: W = - 25.00, *p* = 0.414; 66 mM: W = 4.00, *p* = 0.898; 88 mM: t (10) = 0.958, *p* = 0.361; **Supplementary Figure 1D, H, and L**). Similarly, PAE males exhibited no differences in raw sIPSC amplitude regardless of concentration (44 mM: W = -15.00, *p* = 0.426; 66 mM: W = 3.00, *p* = 0.923; 88 mM: W = 2.00, *p* = 0.966; **Supplementary Figure 1D, H, and L**). Mirroring the raw data, % change from baseline analysis revealed no significant differences in sIPSC amplitude for air (44 mM: t (12) = 0.589, *p* = 0.567; 66 mM: t (10) = 0.650, *p* = 0.531; 88 mM: t (10) = 0.800, *p* = 0.443) or PAE males (44 mM: t (8) = 0.099, *p* = 0.924; 66 mM: t (9) = 0.141, *p* = 0.891; 88 mM: t (10) = 0.038, *p* = 0.970) across all concentrations tested (**Figure 3C**). Lastly, a two-way ANOVA revealed no effects of exposure (F (1, 59) = 0.1502, *p* = 0.700), concentration (F (2, 59) = 0.4235, *p* = 0.657), nor an interaction between the variables (F (2, 59) = 0.3161, *p* = 0.730) on sIPSC amplitude.

#### 3.2.5 G12 PAE shifts sensitivity to acute EtOH in female CeM CRFR1-neurons

As previously described in males, we first analyzed changes to raw sIPSCs. In air females, acute EtOH did not alter raw sIPSC frequency at any concentration tested (44 mM: t (11) = 0.444, *p* = 0.666; 66 mM: t (8) = 0.998, *p* = 0.347; 88 mM: t (10) = 0.093, *p* = 0.928; **Supplementary Figure 2A, E, and I**). Interestingly, in PAE females, we observed a significant decrease in raw sIPSC frequency after 44 mM (W = -70.00, *p* = 0.003) and 66 mM (t (9) = 2.524, *p* = 0.032) application, with no effect at 88 mM (W = 15.00, *p* = 0.426; **Supplementary Figure 2).** When analyzing as % change from baseline, similar to raw sIPSC frequency, no changes were observed in air females (44 mM: t (11) = 1.141, *p* = 0.278; 66 mM: t (8) = 0.501, *p* = 0.630; 88 mM: t (10) = 0.868, *p* = 0.406; **Figure 4A**). A significant decrease in frequency was detected in PAE females at 44 mM (t (11) = 3.805, *p* = 0.003) and 66 mM (t (9) = 2.775, *p* = 0.022), but not at 88 mM (t (8) = 1.068, *p* = 0.317; **Figure 4A**). A two-way ANOVA revealed a main effect of exposure (F (1, 57) = 4.070, *p* = 0.048) and concentration (F (2, 57) = 3.172, *p* = 0.049), but no interaction between the variables (F (2, 57) = 1.709, *p* = 0.190); **Figure 4A**).

**Figure 4.**
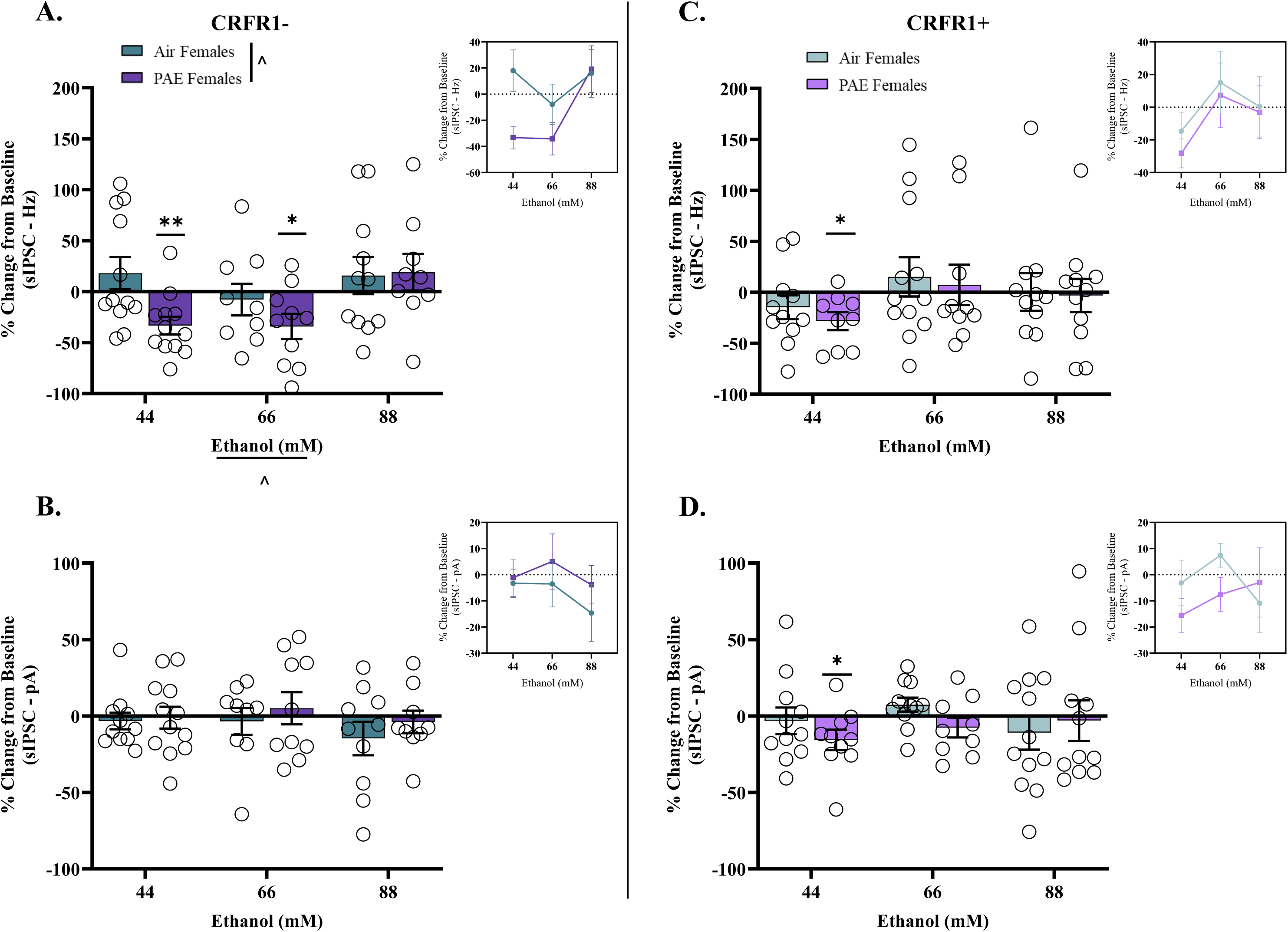
Acute EtOH effects in females represented as % change from baseline. (**A**) A two-way ANOVA revealed a main effects of exposure and concentration in CRFR1- neurons. CRFR1-sIPSC frequency in air females was unaffected by 44 mM (n = 12 cells), 66 mM (n = 9 cells), or 88 mM (n = 11 cells). PAE female CRFR1-sIPSC frequency was significantly decreased by 44 mM (n = 12 cells) and 66 mM (n = 10 cells), but not 88 mM (n = 9 cells). (**B**) No differences were observed for CRFR1-sIPSC amplitude at any concentration for either air or PAE females. (**C**) CRFR1+ sIPSC frequency in air females was unaltered by EtOH at any concentration tested (44 mM: n = 11 cells; 66 mM: n = 12 cells; 88 mM: n = 11 cells). 44 mM significantly decreased CRFR1+ sIPSC frequency in PAE females (n = 9 cells), yet 66 mM (n = 10 cells) or 88 mM (n = 11 cells) had no effect. (**D**) CRFR1+ sIPSC amplitude in PAE females was significantly decreased by 44 mM, but not 66 mM or 88 mM. No significant differences were observed in air females. PAE = prenatal alcohol exposure. CRFR1 = Corticotropin releasing factor receptor type I. sIPSC = spontaneous inhibitory postsynaptic currents. Inset charts represent a different perspective of the same data set. ^ denotes significance from two-way ANOVA. * denotes significance from t-test relative to 0. *p* ≤ 0.05. Bars represent standard error of mean.

**Figure 5.**
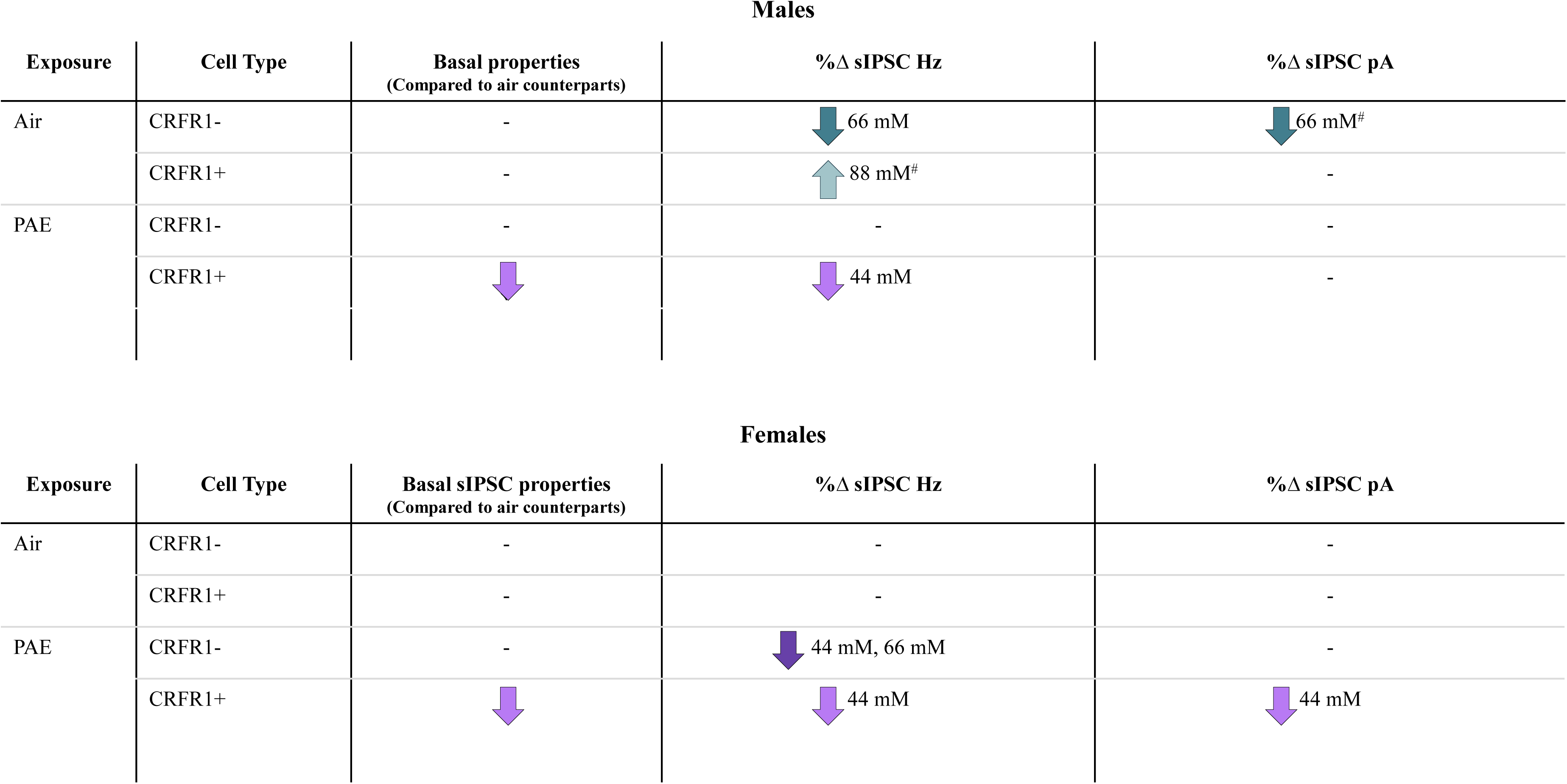
Summary of results. PAE = prenatal alcohol exposure. CRFR1 = Corticotropin releasing factor receptor type I. Hz = frequency. pA = amplitude. sIPSC = spontaneous inhibitory postsynaptic currents. # denotes trends.

Similar to raw sIPSC frequency, raw sIPSC amplitude in air females was insensitive to the effects of acute EtOH at all concentrations tested (44 mM: t (10) = 0.759, *p* = 0.465; 66 mM: t (8) = 0.724, *p* = 0.490; 88 mM: W = -23.00, *p* = 0.275; **Supplementary Figure 2B, F, and J**). Additionally, raw sIPSC amplitude in PAE females was unaffected by EtOH application at all concentrations (44 mM: t (11) = 0.647, *p* = 0.531; 66 mM: W = 5.000, *p* = 0.846; 88 mM: t (8) = 0.473, *p* = 0.649; **Supplementary Figure 2B, F, and J**). Following normalization to % change from baseline, we found no differences in air (44 mM: t (10) = 0.605, *p* = 0.558; 66 mM: t (8) = 0.396, *p* = 0.703; 88 mM: t (9) = 1.339, *p* = 0.214; **Figure 4B**) or PAE females (44 mM: t (11) = 0.158, *p* = 0.878; 66 mM: t (9) = 0.479, *p* = 0.643; 88 mM: t (8) = 0.522, *p* = 0.616; **Figure 4B**) at any concentration tested. Finally, statistical analysis revealed no effect of exposure (F (1, 55) = 1.059, *p* = 0.308), concentration (F (2, 55) = 0.6900, *p* = 0.506), nor an interaction (F (2, 55) = 0.1468, *p* = 0.864; **Figure 4B**).

#### 3.2.6 G12 PAE shifts sensitivity to acute EtOH in female CeM CRFR1+ neurons

Raw sIPSC frequency in air-exposed female CRFR1+ cells was unaffected by acute EtOH at any concentration tested (44 mM: W = -36.00, *p* = 0.123; 66 mM: (W = -14.00, *p* = 0.622); 88 mM: (W = -22.00, *p* = 0.365); **Supplementary Figure 2C, G, and K**). Interestingly, in PAE females, we observed a decrease in raw sIPSC frequency following 44 mM application (W = -41.00, *p* = 0.011), although this was not present at 66 mM (t (9) = 0.956, *p* = 0.364) or 88 mM (W = -4.000, *p* = 0.898). When calculated as a relative change from baseline (%), sIPSC frequency in air females was unaffected by EtOH at all concentrations tested (44 mM: t (10) = 1.258, *p* = 0.237; 66 mM: (t (11) = 0.787, *p* = 0.448); 88 mM: (t (10) = 0.018, *p* = 0.986; **Figure 4B**). Furthermore, 44 mM significantly decreased sIPSC frequency in PAE females (t (8) = 3.196, *p* = 0.013), but 66 mM (t (9) = 0.368, *p* = 0.722) and 88 mM (t (10) = 0.191, *p* = 0.852) had no effect. A two-way ANOVA revealed no effect of exposure (F (1, 58) = 0.3700, *p* = 0.545), concentration (F (2, 58) = 1.897, *p* = 0.159), nor an interaction (F (2, 58) = 0.04584, *p* = 0.955).

Raw sIPSC amplitude in air females did not exhibit any EtOH-induced changes across the range of concentrations tested (44 mM: W = -20.00, *p* = 0.413; 66 mM: t (10) = 1.310, *p* = 0.220; 88 mM: W = -22.00, *p* = 0.424); **Supplementary Figure 2D, H, and L**). Conversely, 44 mM significantly decreased raw sIPSC amplitude in PAE females (t (9) = 2.719, *p* = 0.023), with no changes observed at 66 mM (t (8) = 1.642, *p* = 0.139) or 88 mM (t (10) = 1.208, *p* = 0.255; **Supplementary Figure 2D, H, and L**). Analysis of % change from baseline in air female CRFR1+ cells revealed sIPSC amplitude remained insensitive to acute EtOH, as indicated by the absence of significant changes at any concentration tested (44 mM: t (10) = 0.362, *p* = 0.725; 66 mM: t (10) = 1.626, *p* = 0.135; 88 mM: t (11) = 0.977, *p* = 0.350; **Figure 4D**). However, a significant decrease in PAE females CRFR1+ sIPSC frequency was observed at 44 mM (t (9) = 2.361, *p* = 0.043), with no changes observed at 66 mM (t (8) = 1.179, *p* = 0.272) and 88 mM (t (10) = 0.224, *p* = 0.827; **Figure 4D**). Finally, there was no effect of exposure (F (1, 58) = 0.7332, *p* = 0.395), concentration (F (2, 58) = 0.5147, *p* = 0.600), or an interaction (F (2, 58) = 0.9451, *p* = 0.395) on the dose-response curve.

## 4.0 Discussion

One goal of this study aimed to investigate the effects of *moderate* G12 PAE on basal inhibitory transmission within the CeM of adult rodents. We found that PAE altered postsynaptic responses in CRFR1+ neurons only, as indicated by an increase in sIPSC amplitudes of both males and females, suggesting this cell population exhibits a sensitivity to alcohol *in utero*. Furthermore, we aimed to characterize the effects of acute EtOH on CeM GABAergic transmission and how PAE may disrupt this action in CRFR1+ and CRFR1-neurons in adult CRF_1_-Cre transgenic rats, which to the best of our knowledge is a previously unexplored area of research. Overall, these results demonstrate that PAE differentially modifies acute EtOH’s effects on GABAergic transmission onto CeM CRFR1+ and CRFR1-neurons, in a sex-specific and concentration dependent manner. In males, control animals showed distinct, concentration- and cell type-dependent responses that were blunted or revered in PAE animals. In contrast, control females were unresponsive to EtOH in either cell type, while PAE females exhibit a general inhibition of GABA transmission in both cell types, similar to what we observed in adolescent females (Winchester and Diaz, 2025). These results suggest that PAE shifts EtOH concentration response patterns and cell type interactions which may disrupt neural processing within the CeM, ultimately contributing to alterations in CeM-dependent behaviors. Taken together, these provide meaningful insight into the impact of PAE on neurophysiological responses and the complexity of the relationship between PAE and acute EtOH induced neurotransmission in the CeM.

Our lab has previously reported synaptic modifications associated with PAE in unlabeled CeM neurons of adolescent male (Rouzer and Diaz, 2022) and female (Winchester and Diaz, 2025) Sprague Dawley rats. Specifically, in each instance we observed a decrease in GABA release, reflected by a reduction in sIPSC frequency (Rouzer and Diaz, 2022; Winchester and Diaz, 2025). The current study is the first time our lab has identified synaptic changes in a cell specific manner in the CeM, which revealed an increase in sIPSC amplitude specific to CRFR1+ neurons from both PAE males and females, suggesting post-synaptic modifications as a result of PAE. Collectively, these findings suggest that presynaptic alterations may be transient and return to the level of controls by adulthood, while postsynaptic modifications emerge in adulthood. These postsynaptic modifications could be due to changes in levels of GABA_A_ receptor expression or subunit composition, as seen in other brain regions (i.e. cortex, hippocampus) in different prenatal alcohol animal models (Bailey et al., 2001; Iqbal et al., 2004; Toso et al., 2006). It is equally plausible that a majority of the cells sampled from adolescent rats in our previous research were CRFR1-since we showed that <20% of CeM neurons are positive for CRFR1 mRNA (Rouzer and Diaz, 2022). Importantly, previous research has demonstrated differences in firing properties, and both phasic and tonic inhibitory transmission between CRFR1- and CRFR1+ CeM neurons (Herman et al., 2013). Additionally, the CRF system has been a known target of PAE and alcohol exposure throughout the lifespan (Bertagna et al., 2024; Eisenhardt et al., 2015; Glavas et al., 2007; Kirson et al., 2021; Lam et al., 2018; Patel et al., 2022; Rouzer and Diaz, 2022; Ruffaner-Hanson et al., 2023). Therefore, we have now identified a PAE-induced modification in synaptic transmission specific to CeM CRFR1+ neurons in adults.

Acute EtOH modulation of CeM GABAergic transmission is mediated, in part, by CRFR1 (Nie et al., 2009). It has been shown in different strains and species that acute EtOH (≥ 44 mM) generally potentiates GABA release in the CeM of male rodents (Kirson et al., 2021; Nie et al., 2004; Nie et al., 2009; Roberto et al., 2003; Roberto et al., 2004; Varodayan et al., 2017). Interestingly, we observed an attenuation in GABA release onto CRFR1-neurons of control males (air-exposed) at 66 mM, with no detectable changes in response to 44 or 88 mM. One potential explanation for these differences is the use of a transgenic reporter rodent line in the current study - while advantageous for cell type specificity, insertion of transgenes can alter neurophysiology via insertional mutagenesis, influence and/or disruption of neighboring genes, and promoter promiscuity or leakiness (Matthaei, 2007). While the aforementioned work was performed in wild type animals (i.e. unlabeled neurons), the implementation of reporter lines has allowed researchers to tease apart the impact of acute EtOH on GABA transmission specifically in CRFR1+ neurons. For example, in CRFR1 reporter mice, acute EtOH (44mM) decreased GABA release onto CRFR1+ CeM neurons in naïve adult males (Agoglia et al., 2020). Yet, in the current study, we detected a trend for an increase in GABA release onto CRFR1+ neurons in control males, but only at the highest concentration (88 mM). Even more compelling, PAE males exhibited a decrease in GABA release onto CRFR1+ neurons at the lowest tested concentration (44mM), suggesting that PAE sensitizes this neuronal subtype and produces an opposing effect (reduction versus potentiation of GABA transmission). Despite the noted discrepancies between the current study’s control animals and previous mouse and rat studies, there are known differences in the expression and distribution of CRFR1 in rat vs mouse (Radulovic et al., 1998; Van Pett et al., 2000). Nevertheless, our study is the first to identify cell-specific alterations in response to acute ethanol following PAE in adult male offspring.

EtOH-naïve female rodents are less responsive to the actions of acute EtOH on CeM GABAergic transmission (Agoglia et al., 2020; Agoglia et al., 2022; Kirson et al., 2021; Winchester and Diaz, 2025). When we examined this in a cell-specific manner, air-exposed control females did not exhibit responses to acute EtOH regardless of concentration or cell type, further supporting the existing literature. However, prior EtOH exposure – including PAE or chronic exposure during adulthood - induces sensitivity in these systems (Kirson et al., 2021; Winchester and Diaz, 2025). In adolescent and adult female rats with some history of EtOH exposure, a high concentration of EtOH (88 mM) increased GABA release onto unlabeled neurons (Kirson et al., 2021; Winchester and Diaz, 2025). Interestingly, we found that in PAE females, GABA release was decreased onto CRFR1+ and CRFR1-neurons at the lowest concentration (44 mM) and CRFR1-at the middle concentration (66 mM). Typically, in naïve, control or alcohol-dependent male rodents, CRF and/or acute EtOH produce similar neurophysiological responses in CeM neurons (i.e. GABA potentiation). Acute EtOH’s potentiating effect on GABA transmission has been shown to be mediated, in part, through CRF and/or CRFR1 (Nie et al., 2009). While adult naïve female Sprague Dawley rats have been shown to be insensitive to different concentrations of CRF (Rodriguez et al., 2022), we previously found that the CRFR1 selective agonist, Stressin-1, significantly attenuated GABA release onto unlabeled neurons of naïve female Sprague Dawley rats, with adult females showing a higher sensitivity to lower concentrations of Stressin-1 (Rouzer and Diaz, 2021). Additionally, PAE blunted the effect of Stressin-1 on GABA transmission in adolescent females (Rouzer and Diaz, 2022). The findings from the current study extend our understanding on the impact of *in utero* EtOH exposure on the CRF/CRFR1 CeM system in females, showing that PAE unmasks an effect of acute EtOH in CRFR1- and CRFR1+ neurons, favoring a reduction in GABAergic signaling in adulthood. Importantly, this suggests that the established interactions between acute EtOH and CRF/CRFR1 in males differ in females. Additionally, these findings suggest that with age, the system becomes more sensitive to EtOH, with lower concentrations in adulthood producing opposite effects to those observed in adolescence at higher concentrations.

Notably, a portion of our findings diverge from previous reports examining acute EtOH effects in the CeM of male rodents. Several methodological differences may account for the discrepancies. For example, cutting/slicing solution and temperature can alter the frequency of sIPSCs (Avegno et al., 2019). Specific to the CeM, researchers have reported changes in neuronal excitability — such as hyperpolarization of resting membrane potential — increased sIPSC frequency, and enhanced drug effects when recordings were performed under physiologically relevant temperature as opposed to room temperature (Avegno et al., 2019). Furthermore, the use of NMDG as a cutting solution significantly increased basal sIPSC frequency (Avegno et al., 2019).

Additionally, differences in synaptic blockers to isolate GABAergic transmission can contribute to discrepancies across laboratories. For example, quinoxaline derivatives like DNQX and CNQX have been shown to produce membrane depolarizations in rat thalamic and hippocampal neurons (Lee et al., 2010; Maccaferri and Dingledine, 2002). Collectively, these studies highlight how procedural differences influence neurophysiological measures, such as those examined in this and other comparable studies, which may contribute to the discrepancies between the current study and previous work. Furthermore, the CeA is known to express a variety of neurotransmitters and neuropeptide systems (O’Leary et al., 2022; Wang et al., 2023; Yeh et al., 2024) which may contribute to the variability in responsiveness to acute EtOH. Previous analysis of a dose response curve (11-66 mM) revealed ∼30% of neurons were reported to be non-responders or showed decreased GABAergic transmission in response to 44 mM EtOH in the CeA (Roberto et al., 2003). Therefore, it is also possible that removal of non-potentiated or non-responding neurons from analyses could account for the variable findings across studies. The current study included all recorded CRFR1+ and CRFR1-neurons in our analyses, unless a cell statistically qualified as an outlier. While selective inclusion can be valuable for mechanistic dissection, we felt that inclusion of the full heterogeneity of EtOH effects in a highly heterogeneous nucleus was important.

While reporter lines can provide valuable insight into the contribution of specific neuronal subtypes or circuitry involved in different disease models, there are still limitations to using these models, some of which have been previously outlined. Examination of multiple CRF-reporter mice models has revealed incongruence of the reporter gene compared to the endogenous gene, where expression may have strong overlap in some brain region but little to none in others (Chen et al., 2015). Furthermore, once reporter genes are activated, they are continually expressed (Deussing and Chen, 2018), therefore some CRFR1+ neurons recorded in our study may have only transiently expressed the receptor, and not necessarily at the time of recording. Nevertheless, the current study extends the field’s knowledge regarding the long-term effects of *moderate* PAE on synaptic function and the predisposition to altered responses to acute EtOH in adult offspring. Furthermore, this study highlights the intricacies and complexities of the CeM CRFR1 system and reaffirms that the CeM CRFR1 is a major target of PAE.

## Acknowledgements

This study was funded by NIAAA grants R01AA028566, R01AA028566 supplement, F31AA032198, and P50AA017823.

---

**Supplementary figure 1.** Raw sIPSC frequency and amplitudes for CRFR1- & CRFR1+ in males. PAE = prenatal alcohol exposure. CRFR1 = Corticotropin releasing factor receptor type I. sIPSC = spontaneous inhibitory postsynaptic currents. * denotes significance from paired t-test. *p* ≤ 0.05. Bars represent standard error of mean.

**Supplementary figure 2.** Raw sIPSC frequency and amplitudes for CRFR1- & CRFR1+ in females. PAE = prenatal alcohol exposure. CRFR1 = Corticotropin releasing factor receptor type I. sIPSC = spontaneous inhibitory postsynaptic currents. * denotes significance from paired t-test. *p* ≤ 0.05. Bars represent standard error of mean.

