## Supplementary figures and images for "Moderate prenatal alcohol exposure differentially alters acute ethanol sensitivity of GABAergic transmission in CRFR1- and CRFR1+ CeM Neurons"

### Supplemental figures

**Supplementary Figure 1.** Males: Raw sIPSC.

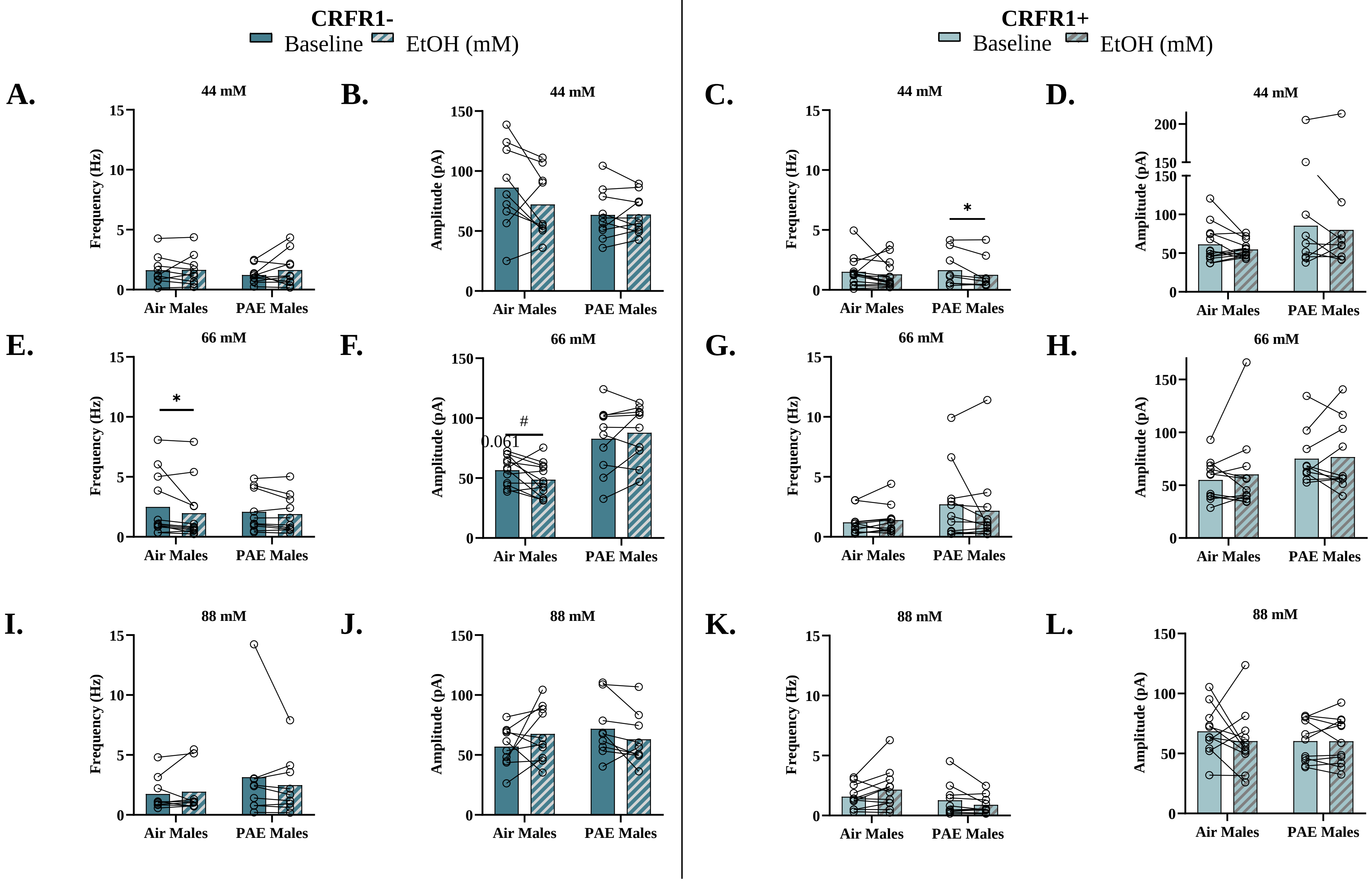

**Supplementary Figure 2.** Females: Raw sIPSC.

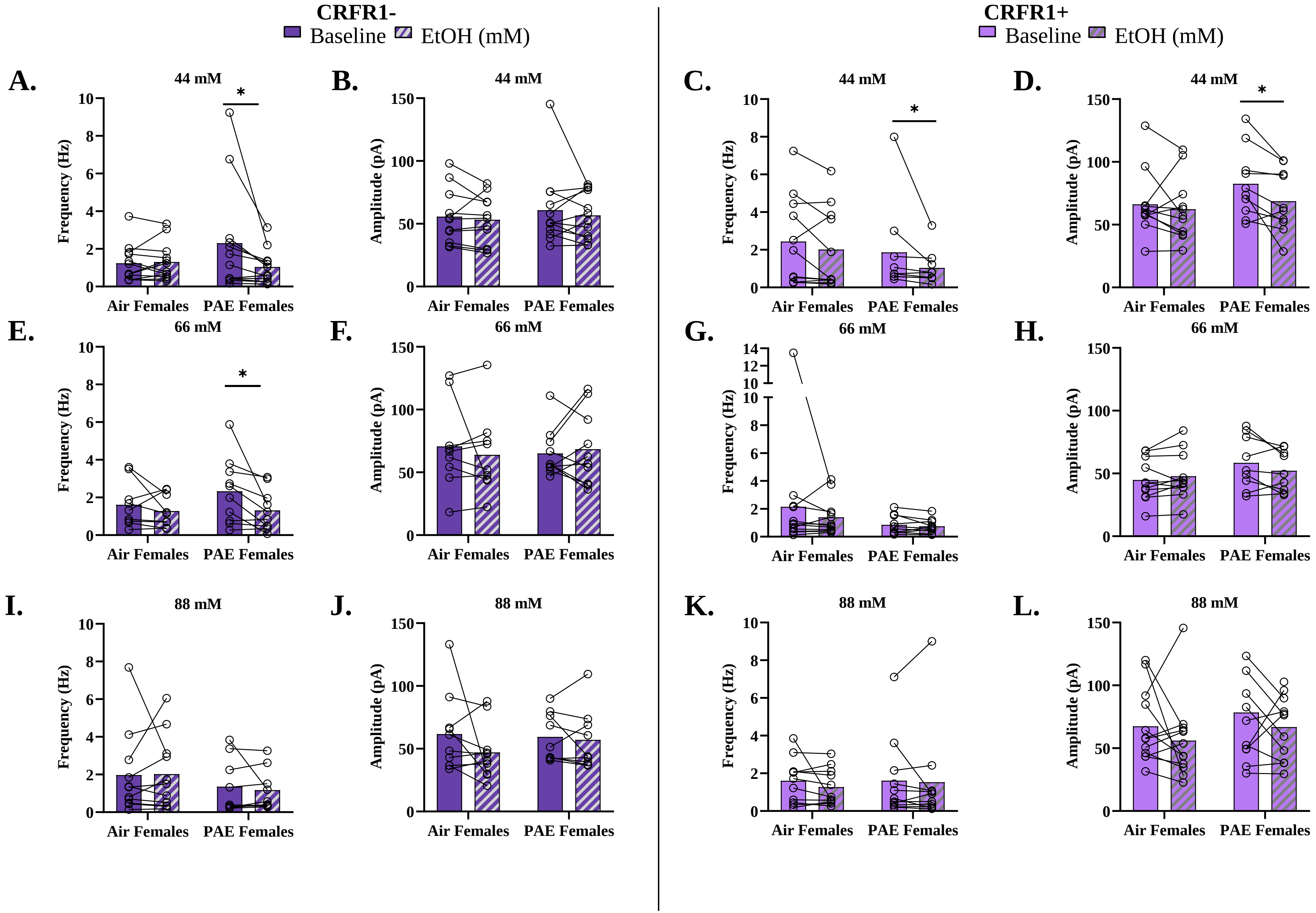
